## Supplemental Figures for "The C-terminus of the multi-drug efflux pump EmrE prevents proton leak by gating transport"

**This PDF file includes:**

SI Figures 3.1 to 8.4

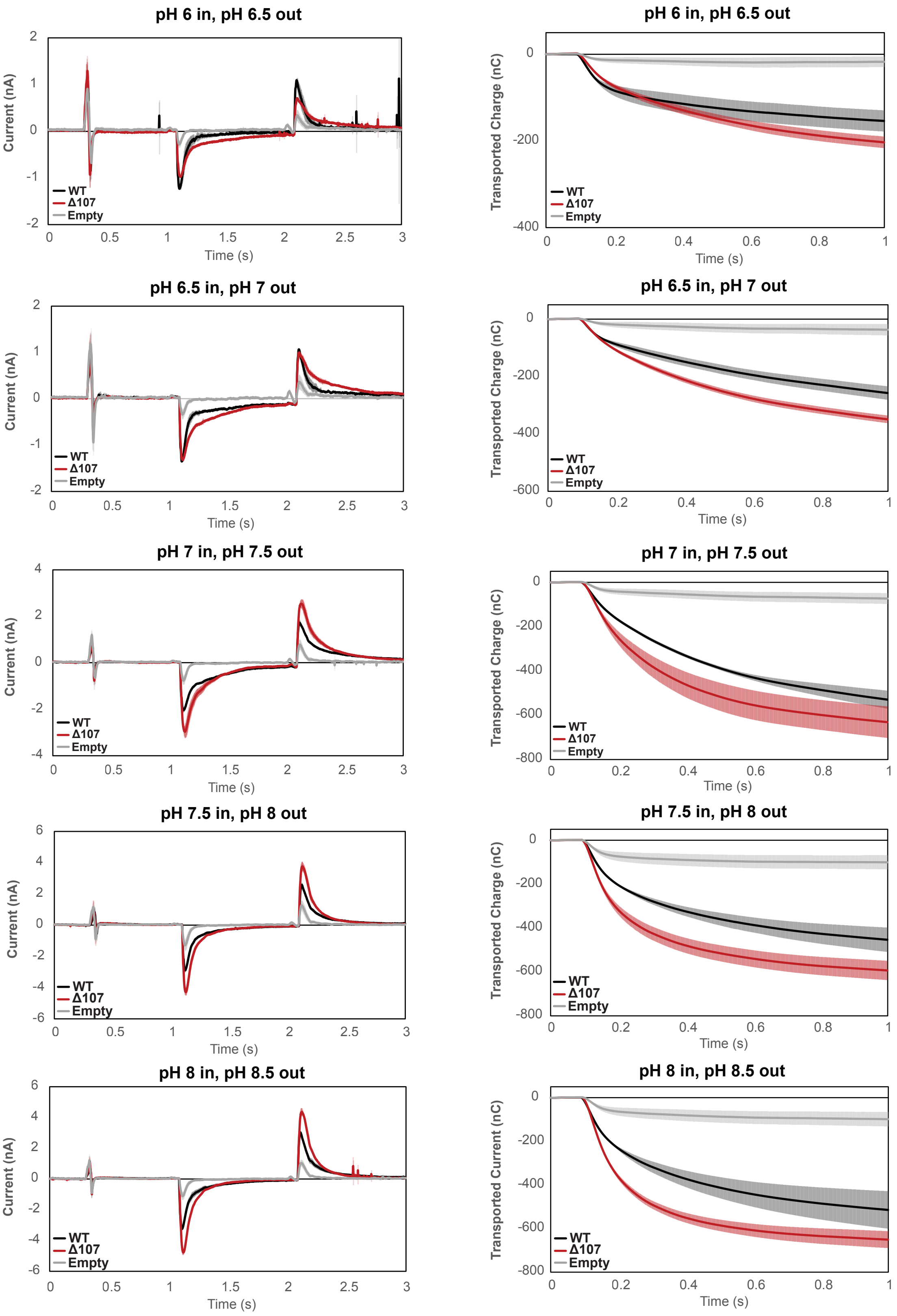


**Figure 3.1. Averaged currents and integrated transport curves of WT-EmrE, ∆107-EmrE, and empty liposomes in the presence of different pH gradients.** Graphs represent the current upon formation of a pH gradient and then return to starting conditions (left) and integrated current upon formation of the pH gradient (right). The magnitude of the pH gradient is constant, but the absolute pH differs (top to bottom). Each curve is an average of 3 technical replicates of individually prepared sensors and error bars represent the standard deviation from the mean. The kinetics of uncoupled-proton leak between WT- EmrE and ∆107-EmrE is distinct at different absolute pH, but the overall transported charge by ∆107-EmrE is consistently higher.

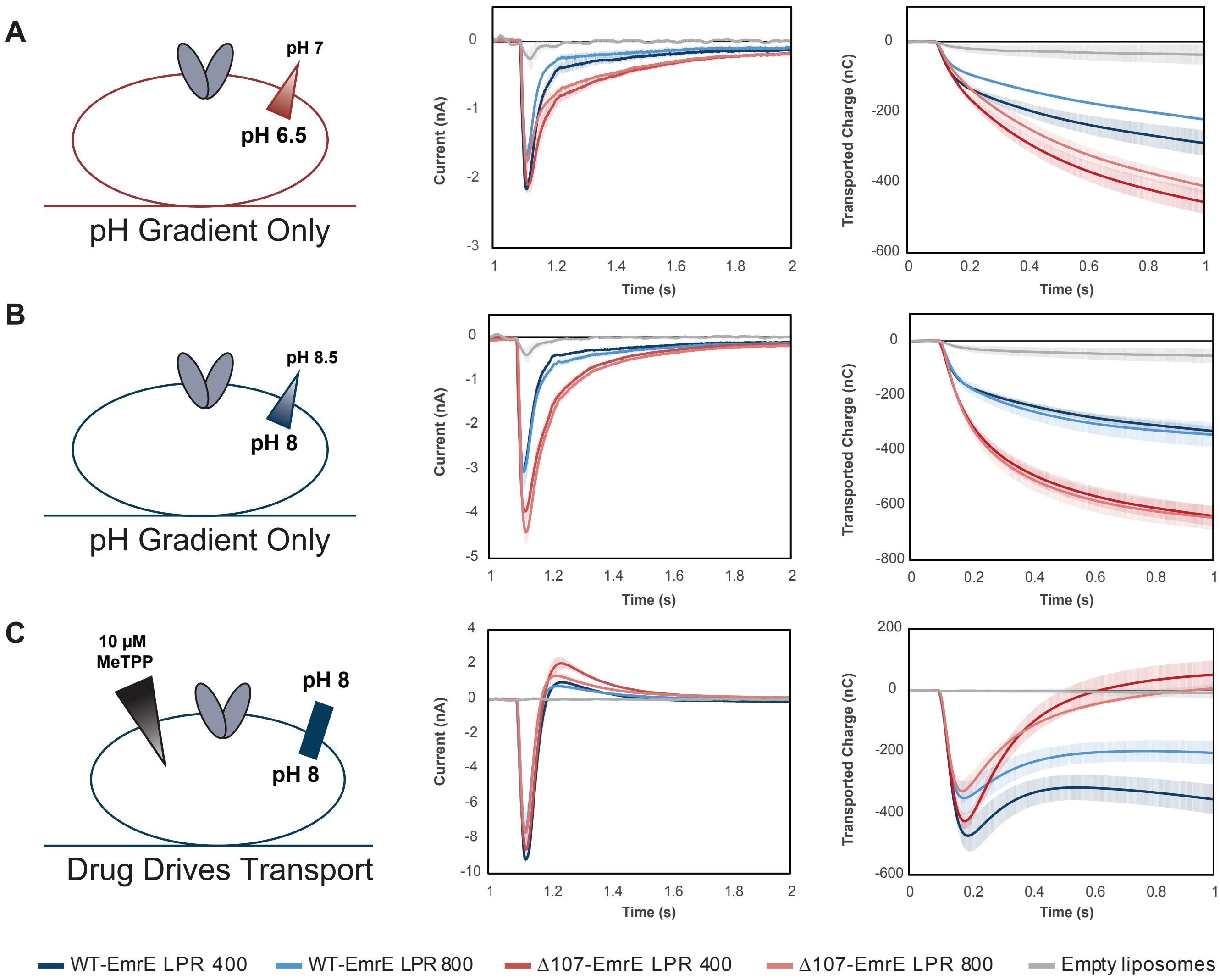


**Figure 3.2. Averaged currents and integrated transport curves of WT-EmrE and Δ107-EmrE with different Lipid to Protein Ratios** (LPRs). Comparing more than one LPR will alter the kinetics, but not the thermodynamics of transport. As such, comparing the peak currents for more than one LPR for a given set of gradient conditions allows us to ensure the signal is dominated by steady state transport rather than pre-steady state electrogenic partial reactions, such as substrate binding or proton release from the transporter. (A) The peak currents for WT-EmrE with different LPR do not match and the peak current for Δ107-EmrE with different LPR do not match, indicating contributions from a pre-steady state process (left). Instead, the LPR 400 data for WT- and Δ107-EmrE peak currents match and the LPR800 data for WT- and Δ107-EmrE match, suggesting that release of protons from the transporter from E14 when the external buffer is switched dominates the peak current in this pH range. No substrate is present and H110 is only present in one construct, so substrate binding or proton release from H110 are unlikely to contribute significantly to this signal. However, the slower kinetics of the off rates for these currents are not consistent with a strictly pre-steady state process, indicating transport of protons, or leak, is also occurring for both constructs consistent with the initial jump in fluorescence followed by a slower leak over time that was seen in the pyranine assay. The similarity of the integrated currents (transported charge over time, right) for the different LPRs of each construct and the consistently higher signal in Δ107-EmrE across LPRs further signifies the predominance of uncoupled proton leak upon deletion of the C-terminal tail. If pre-steady state signals were dominant, we would expect WT-EmrE to have a higher signal due to the contribution of an additional titratable residue (H110). (B) With the same magnitude gradient at high pH, the transporters will be mostly de-protonated, such that we do not see peak currents that are dependent on the density of protein as in A. Instead, the peak currents and off-rates are dependent on the presence or absence of the tail (left) and the transported charge over time is the same for both LPRs of each construct (right). (C) In the absence of a pH gradient, addition of an infinite drug gradient to the outside of the liposomes containing de-protonated

transporters reveals how coupled transport impacts the observed current. The infinite inward-directed drug gradient causes rapid coupled antiport of two protons out of the liposome for every one $MeTPP^+$ molecule transported in (initial negative peak current). This creates both a negative inside potential that inhibits further transport and increases the external proton concentration. The resulting inward-directed proton gradient and negative-inside potential drives back transport of protons into the liposome (slower phase, positive current). This positive current is greater for ∆107-EmrE than for WT-EmrE, with the net transported charge returning to near baseline for ∆107-EmrE. There is only limited proton backflow through WT-EmrE, which maintains tighter coupling due to the C-terminal tail. Each curve is an average of at least 3 technical replicates of individually prepared sensors and error bars represent the standard deviation from the mean.

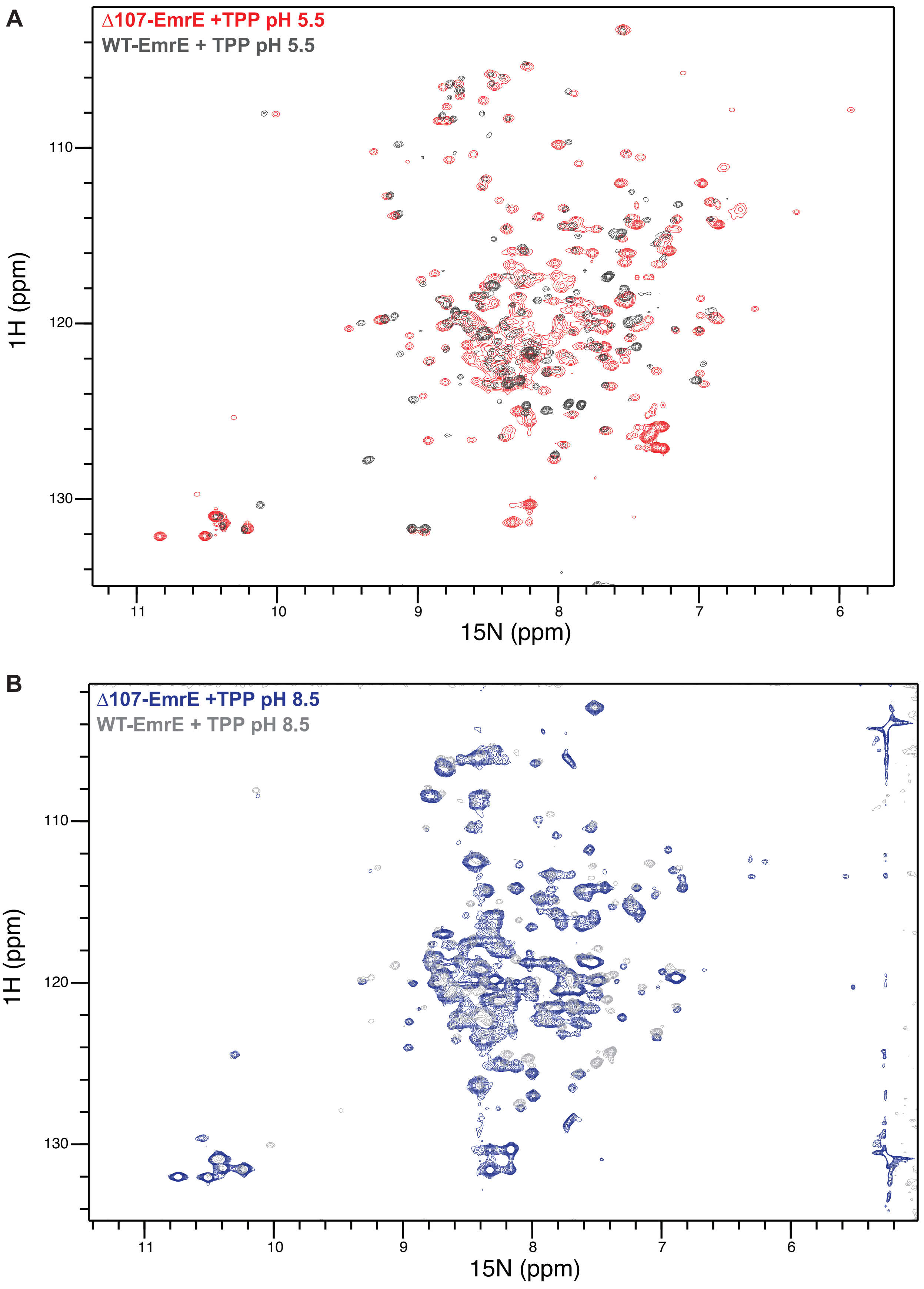


**Figure 4.1. Overlay of drug bound WT- and Δ107-EmrE at low and high pH.** 1H-15N TROSY-HSQC spectra of WT- and Δ107-EmrE are at pH 5.5 (A) and pH 8.5 (B) are highly similar suggesting the C-terminally truncated Δ107-EmrE mutant is properly folded and has an intact binding site.

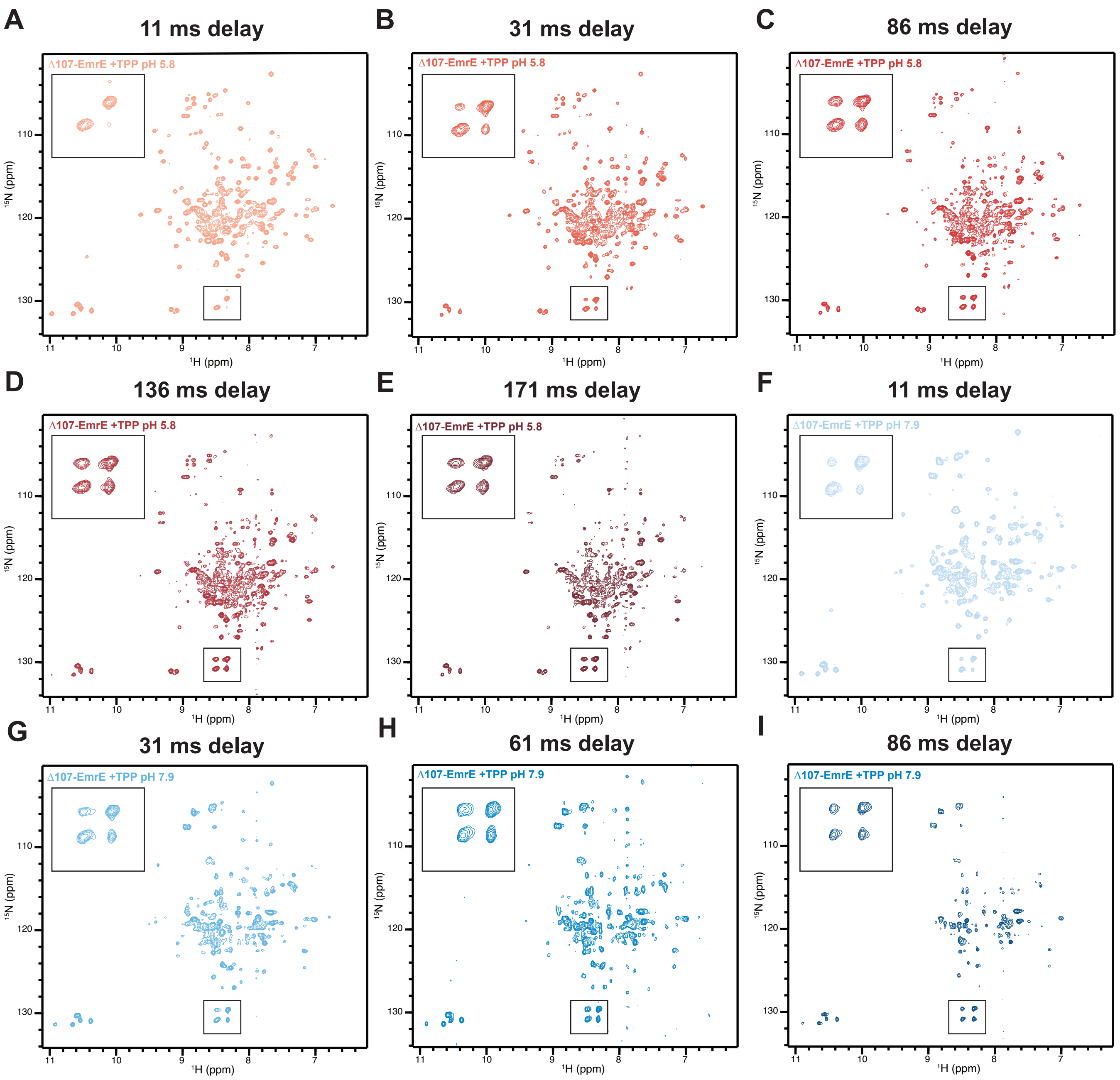


**Figure 4.2. Full ZZ-exchange TROSY-HSQC spectra of the low and high pH Δ107-EmrE with TPP+.** Spectra shown were collected with the indicated delay time between recording the 15N and 1H chemical shifts. The indicated boxes were enlarged to highlight the increasing intensity of the cross-peaks compared to the auto-peaks of R106, the new C-terminus of the truncated construct.

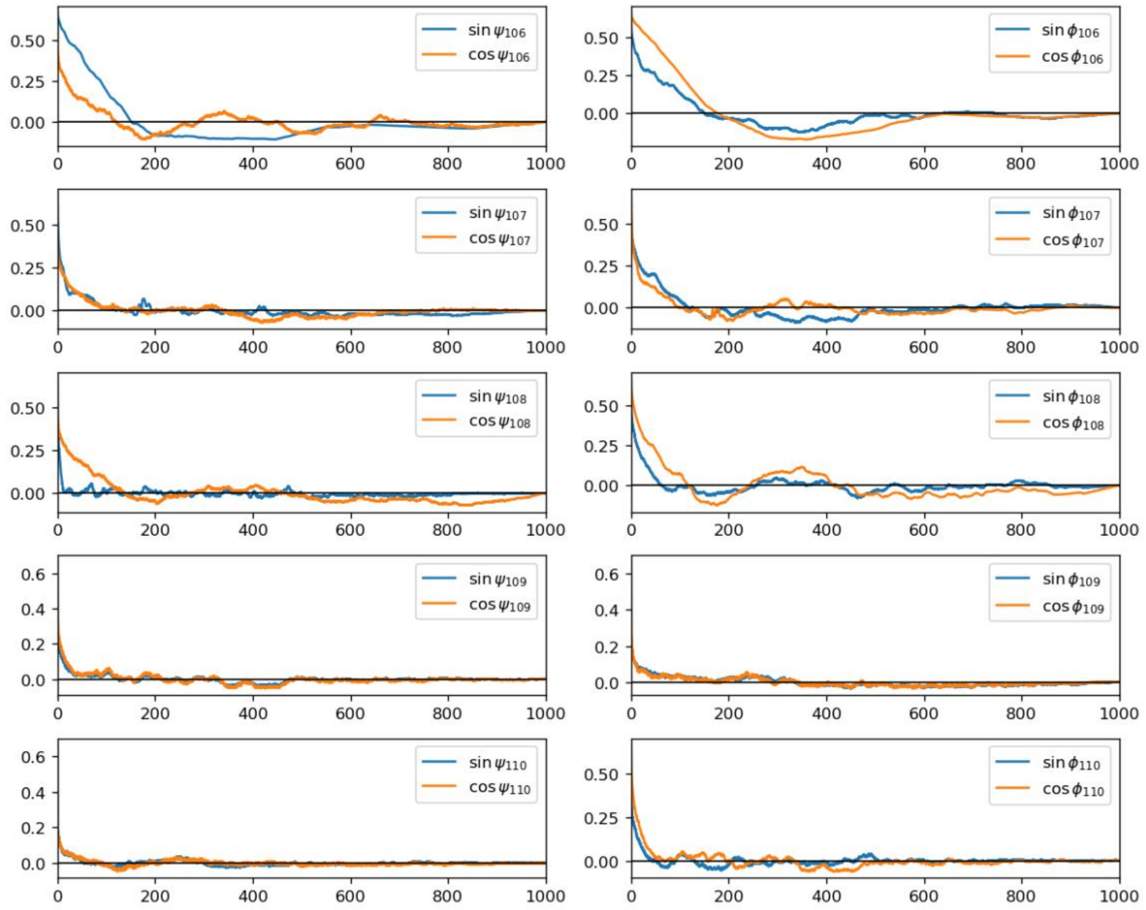


**Figure 5.1 Autocorrelation function of dihedral angles of the tail.** The normalized autocorrelation function of psi- and phi- dihedral angle of the C-terminal tail backbone was plotted against time (unit: ns), averaged over three replicas. This shows the equilibration of the tail is fast except for residue 106 because of a potential salt bridge (with D84) and its proximity to helix 4, which has a stable secondary structure and a rigid backbone. The fast equilibration allows us to conclude the sampling of the tail structure is close to ergodic.

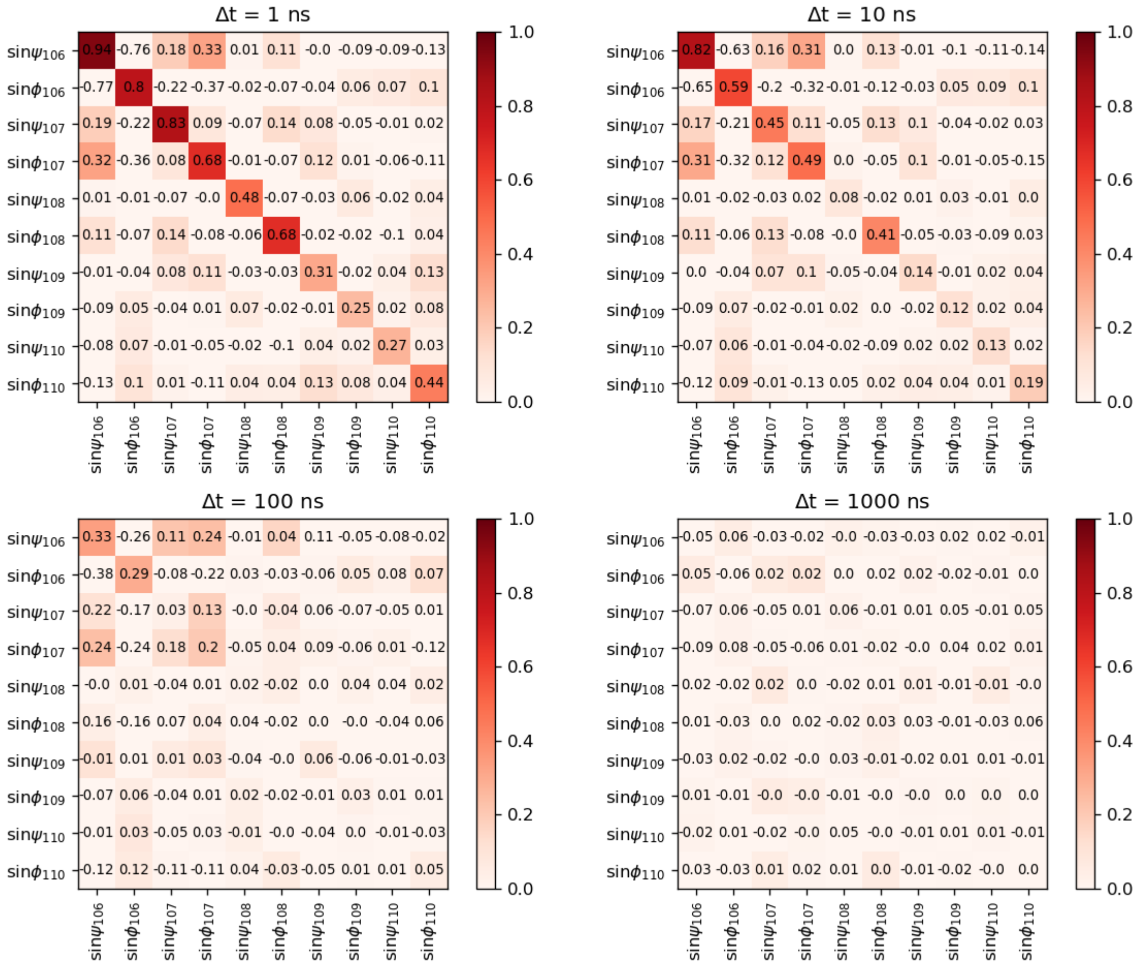


**Figure 5.2 Cross-correlation function of dihedral angles of the tail at four given lag times.** The normalized cross-correlation function of psi- and phi- dihedral angle of the C-terminal tail backbone at four different timescales. Cross-correlation is minimal except very close angles. The fast equilibration allows us to conclude the sampling of the tail structure is close to ergodic.

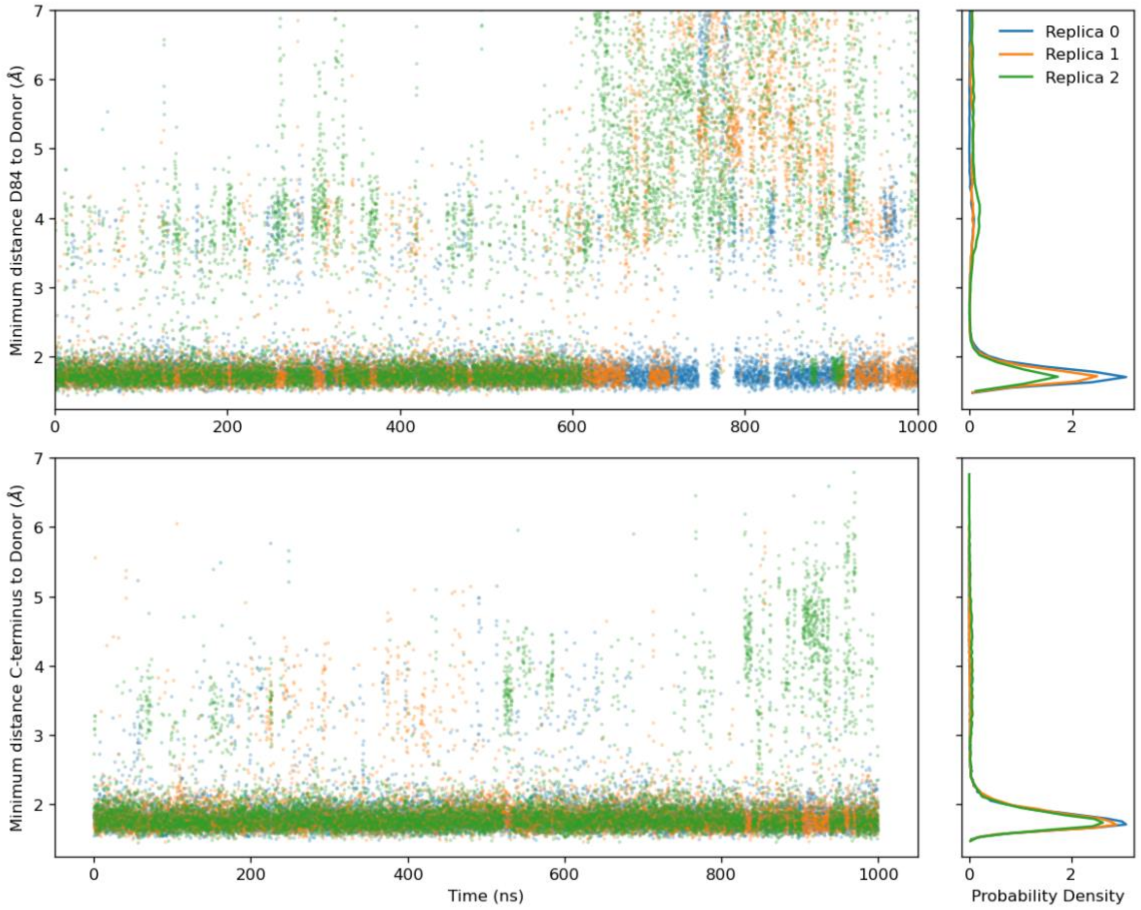


**Figure 5.3. Hydrogen bonding of D84 and the C-terminal tail.** The hydrogen bonding of D84 and the carboxylate group of the C-terminal can be characterized by the distances between these oxygens and their nearest donor hydrogen. The left two panels show the time series of these distances, and the right two panels shows the probability density. The donor for the C-terminus is the side chain of T56. The donor for D84 is primarily R106, but there is a small probability density of hydrogen bonding to S105 near 4 Å.

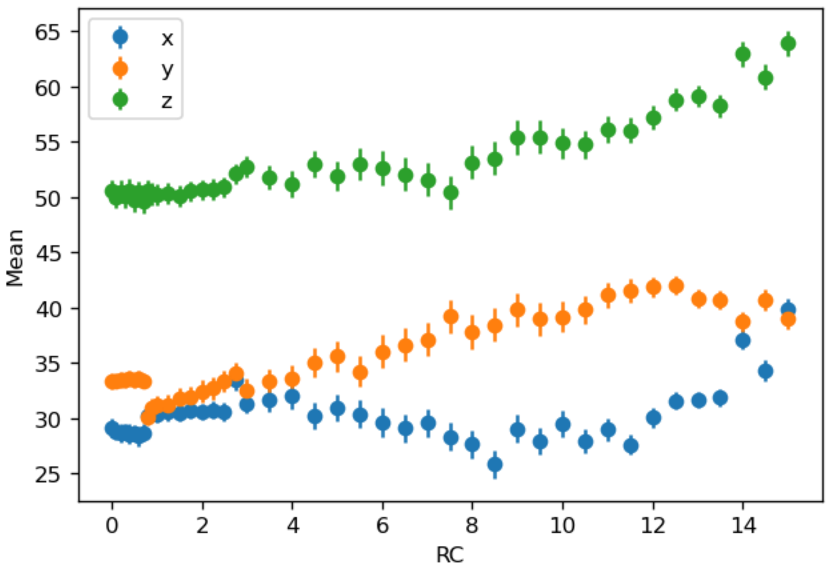


**Figure 6.1. Average Cartesian coordinates of umbrella windows.** The cartesian coordinates of each umbrella window by reaction coordinate (RC) along with the standard deviation is plotted. This shows the continuity of the motion of the CEC when biasing the projected CV. The jump at x~1 Angstrom corresponds to an E14 sidechain rotation event.

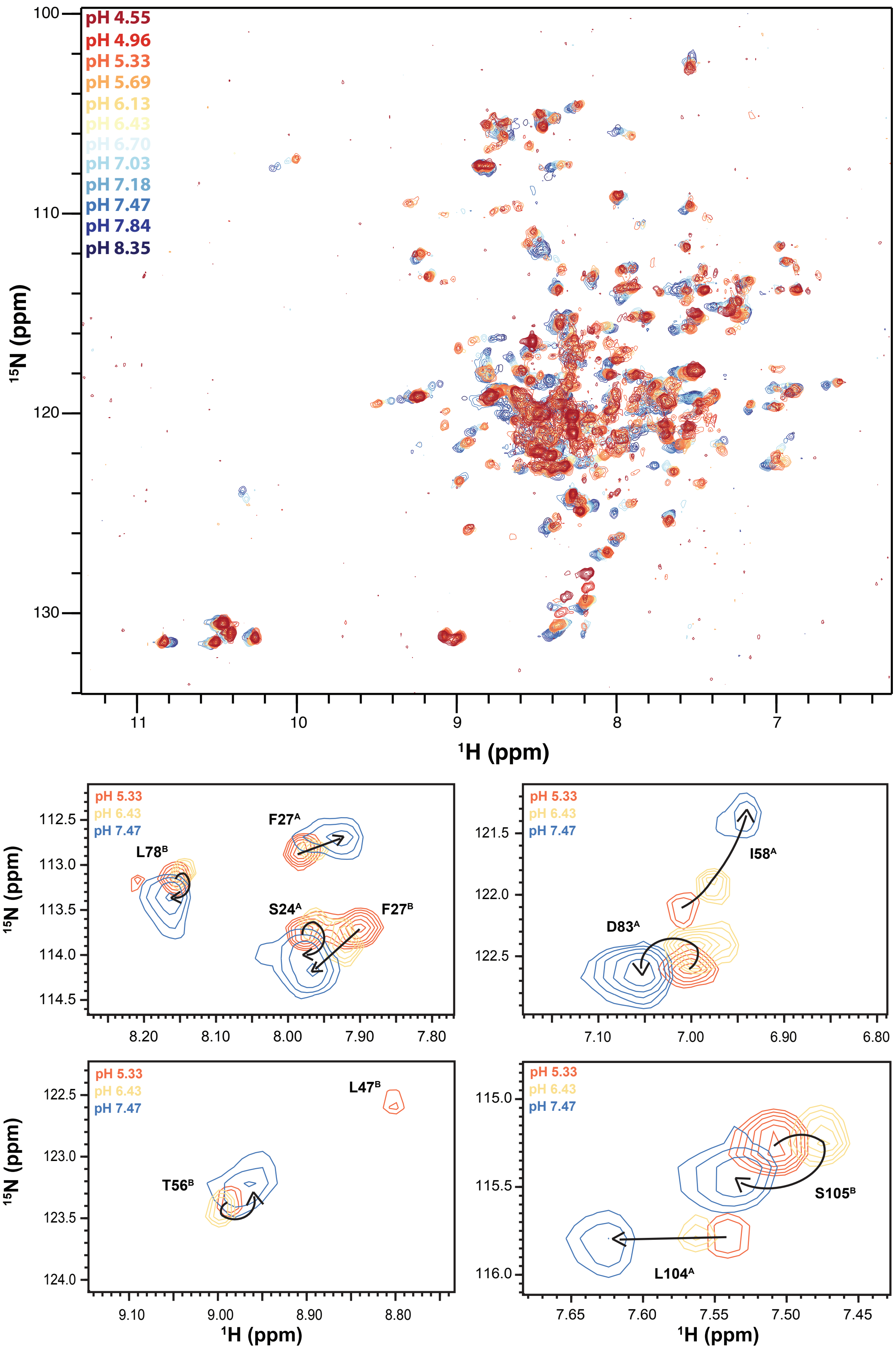


**Figure 7.1. NMR pH titration of TPP-bound Δ107-EmrE.** The pH titration of TPP-bound Δ107-EmrE in isotropic bicelles was performed at 45 C and individual spectra were collected by mixing portions of a high pH and low pH sample in equivalent buffers to obtain the given pH and collecting 1H-15N TROSY HSQCs. While it appears the majority of titrating peaks are moving along a straight line, indicative of one protonation event, peaks can be seen that display curved titration paths and large jumps between pH points.

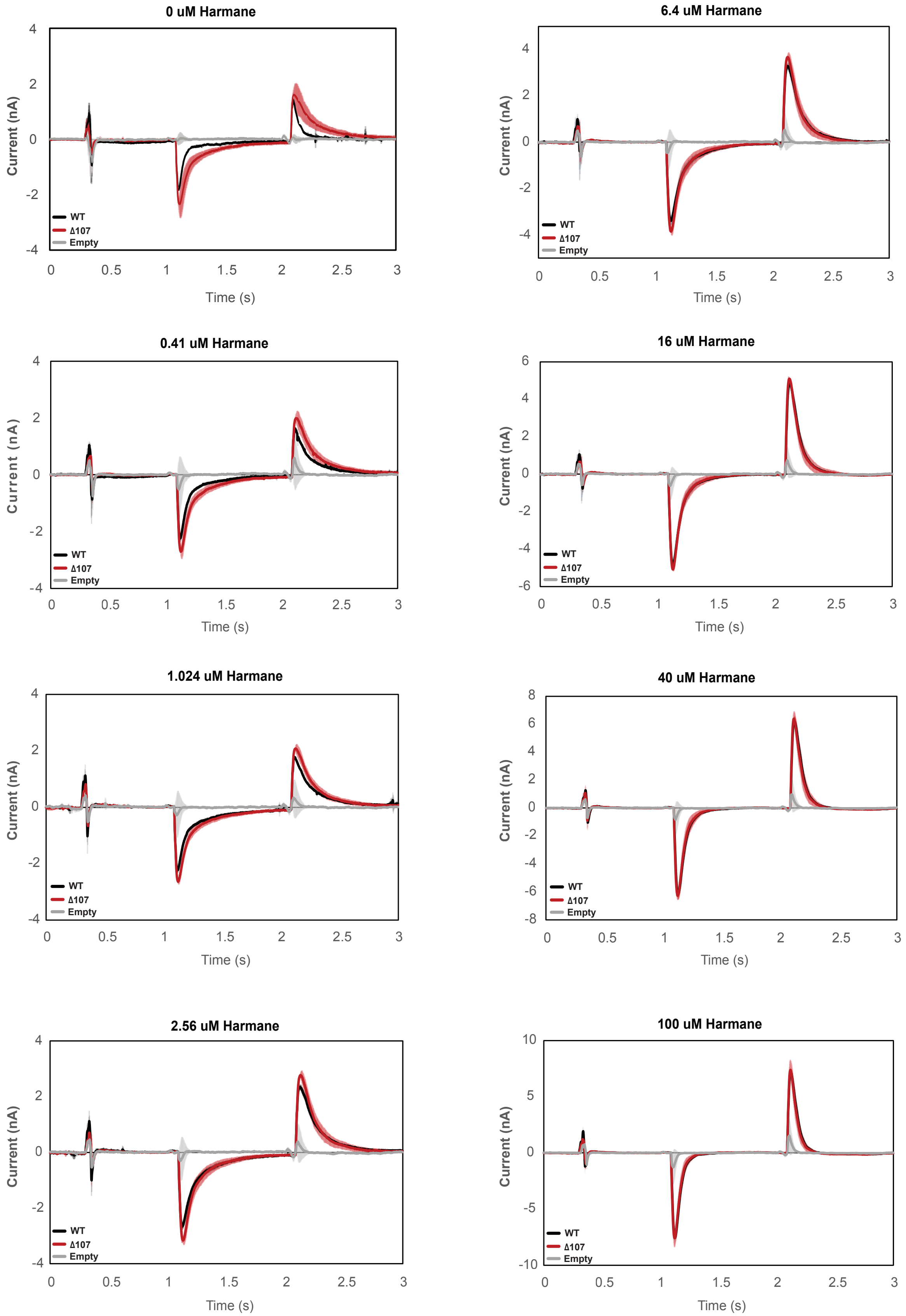


**Figure 8.1. Averaged currents of WT-EmrE,** Δ107-EmrE, and empty liposomes in the presence of different concentrations of harmane. Graphs represent the current under a constant pH (pH 6.5 inside vs pH 7 outside) in the presence of varying harmane concentrations. Each curve is an average of 3 technical replicates of individually prepared sensors and error bars are the standard deviation from the mean. The initial difference in the amount of uncoupled-proton leak between WT- EmrE and Δ107-EmrE is abolished in the presence of higher concentrations of harmane.

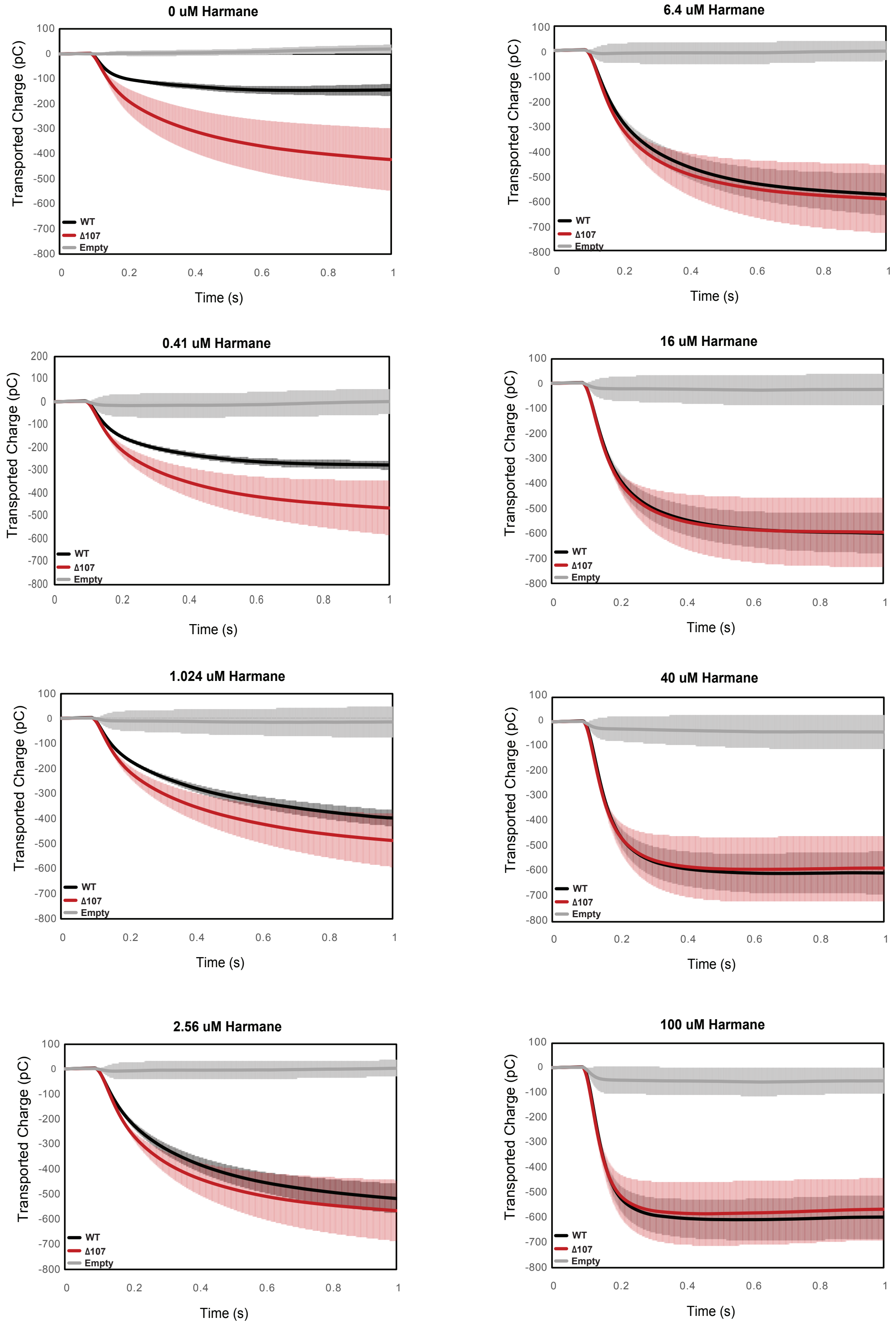


**Figure 8.2. Integrated transport curves of WT-EmrE, ∆107-EmrE, and empty liposomes in the presence of different concentrations of harmane.** Graphs represent the transported charge under a constant pH (pH 6.5 inside vs pH 7 outside) in the presence of varying harmane concentrations. Each curve is an average of 3 technical replicates of individually prepared sensors and error bars are the standard deviation from the mean. The initial difference in the amount of uncoupled-proton leak between WT- EmrE and ∆107-EmrE is abolished in the presence of higher concentrations of harmane.

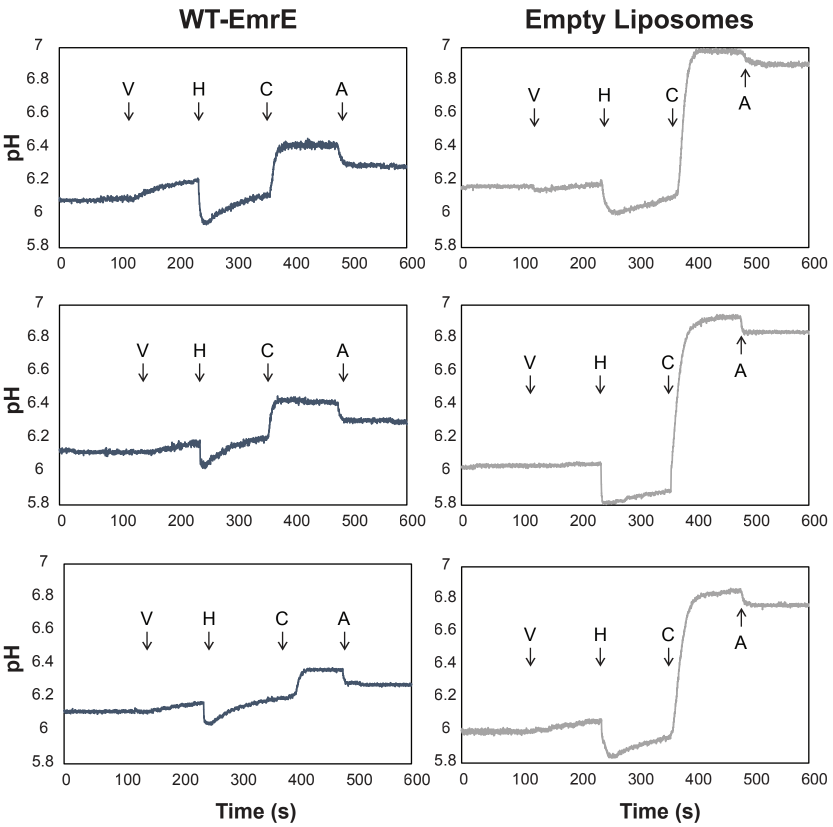


**Figure 8.3. pH detected liposomal leak assay shows harmane dissipates ∆pH in an EmrE-dependent manner.** Proton-tight proteoliposomes and empty liposomes with a 3:1 ratio of POPC:POPG were prepared in a pH 7 internal buffer (50 mM MOPS, 100 mM KCl) and buffer exchanged into a pH 6 external buffer (50 µM MES, 1 mM KCl, and 99 mM NaCl) such that a ∆pH and ∆ are present. The basal pH-dependent leak of WT-EmrE leads to a difference of ~0.1 pH units for WT-EmrE containing proteoliposomes relative to empty liposomes in the time it takes to buffer exchange and commence the measurement, however the flat line prior to ionophore addition reveals this leak is slow on the timescale of the assay. Addition of valinomycin (V) allows a small number of protons to be exchanged across the membrane for both empty liposomes and proteoliposomes. Addition of harmane (H) lowers the pH of the weakly buffered solution external to the liposomes, then induces an EmrE-dependent, logarithmic leak in proteoliposomes that is distinct from the slow, linear leak present in empty liposomes. This leak dissipates the ∆pH across the proteoliposomes to a much greater extent than for the empty liposomes such that addition of the ionophore CCCP (C) has a greatly reduced effect. Finally, a set amount of hydrochloric acid (A) is added for scale.

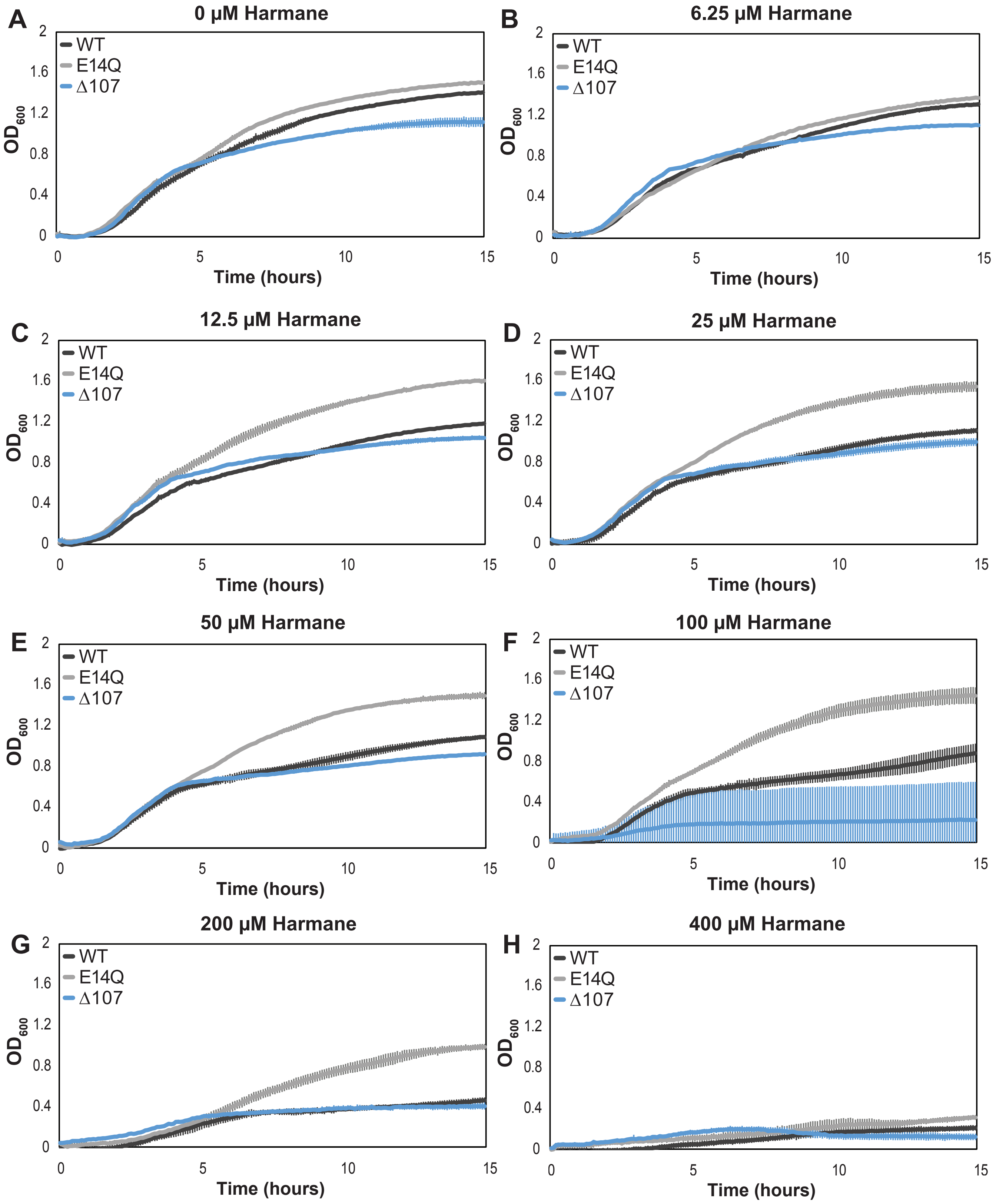


**Figure 8.4. Growth assays show differential impact of harmane on growth of *E. coli* expressing WT-, E14Q-, or Δ107-EmrE.** Graphs represent the average growth in the presence of varying harmane concentrations as measured by OD600. Each curve is representative of an average of six technical replicates stemming from two biological replicates and error bars are the standard deviation from the mean. Again, the growth defect in Δ107-EmrE expressing cells is evident in (A) and while the E14Q-EmrE expressing cells do not appear to have grown as well as is typical in (B), overall, the convergence of WT- and Δ107-EmrE expressing cells as harmane concentration increases can be seen (C-E) before all cells begin to die due to a non-specific toxicity (F-H).
